# Extra-cellular Matrix in cell aggregates is a proxy to mechanically control cell proliferation and motility

**DOI:** 10.1101/2020.10.06.328252

**Authors:** Monika E. Dolega, Sylvain Monnier, Benjamin Brunel, Jean-François Joanny, Pierre Recho, Giovanni Cappello

## Abstract

Imposed deformations play an important role in morphogenesis and tissue homeostasis, both in normal and pathological conditions. To perceive mechanical perturbations of different types and magnitudes, tissues need appropriate detectors, with a compliance that matches the perturbation amplitude. By comparing results of selective osmotic compressions of cells within multicellular aggregates with small osmolites and global aggregate compressions with big osmolites, we show that global compressions have a strong impact on the aggregates growth and internal cell motility, while selective compressions of same magnitude have almost no effect. Both compressions alter the volume of individual cells in the same way but, by draining the water out of the extracellular matrix, the global one imposes a residual compressive mechanical stress on the cells while the selective one does not. We conclude that, in aggregates, the extracellular matrix is as a sensor which mechanically regulates cell proliferation and migration in a 3D environment.

## 1 Introduction

Aside from biochemical signaling, cellular function and fate also depend on the mechanical state of the surrounding extracellular matrix (ECM) ***(Humphrey et al., 2014***). The ECM is a non-cellular component of tissues providing a scaffold for cellular adhesion and triggering numerous mechanotransduction pathways, necessary for morphogenesis and homeostasis ***(Vogel, 2018***). An increasing number of studies *in vivo* and *in vitro* shows that changing the mechanical properties of the ECM by reimplanting tissues or changing the stiffness of the adherent substrate is suffcient to reverse aging ***(Segel et al., 2019***), accelerate developmental processes ***(Barriga et al., 2018)*** or reverse the pathological state in cancer ***(Paszek et al., 2005; Tanner et al., 2012***).

The importance of the mechanical context in cancer has for long been highlighted by experiments altering the composition and stiffness of ECM ***(Levental et al., 2009***), but more recently it has also been shown that the growth of tumors is modulated by the accumulation of the mechanical pressure caused by the hyper-proliferation of cells during tumor expansion in a confined environment ***(Fernandez-Sanchez et al., 2010; Nia et al., 2016***). Such physiological growth under pressure has been studied also *in vitro* in the multicellular context. When multicellular aggregates are confined by soft gels (***Helmlinger et al., 1997***; ***Alessandri et al., 2013***; ***Taubenberger et al., 2019***) or submitted to a gentle osmotic compression (***Montel et al., 2011***; ***Dolega et al., 2017***), their growth is substantially reduced. It has been demonstrated that the cell cytoskeleton is involved in the response to compression and can trigger the growth impediment through cycle-cell inhibition (***Taubenberger et al., 2019***; ***Delarue et al., 2014***). In addition, cellular volume has been recently proposed to be a key parameter in the mechanosensitive pathway (***Delarue et al., 2014***; ***Han et al., 2020***). Nevertheless, it is not known how such mild global compression is transduced to the individual cells of the aggregate to alter their proliferation.

Here, we introduce the hypothesis that cells mainly respond to the mechanical stress transmitted by the ECM, when the aggregate is under compression. The hypothesis is motivated by two evidences. First, aggregates are a composite material made of cells, extracellular matrix and interstitial fluid. The ECM being a a soft device 100 to 1000-fold more compressible than the cells, it absorbs most of the deformation, but still transmits the mechanical stress to the cells. Second, whereas an osmotic pressure of few kPa strongly reduces cell proliferation inside multicellular aggregates, an identical pressure has no effect on individual cells cultured on a Petri dish, in the absence of ECM.

To test the hypothesis that cells respond to the ECM deformation, we introduce an experimental method that uncouples the cell volume change from the mechanical stress transmitted to the cells through the ECM. We apply this method both for both multicellular aggregates and individual cells embedded in a gelified ECM. In parallel, we present a theoretical framework to estimate both the displacement and the stress at the ECM/cell interface in response of an osmotic compression, and verify experimentally its qualitative prediction. At longer timescale, we probe the effect of the ECM compression on the cellular response, in terms of proliferation and motility. Our results show that, even in the absence of cell deformation, the ECM deformation alone is an important factor determining these properties.

## 2 Results

### 2.1 Selective-compression method

We developed a simple method to either selectively compress cells embedded in ECM or the whole complex composed of ECM and cells. This method is based on the use of osmolytes of different sizes. When large enough, the osmolytes do not infiltrate the ECM and thus compress the whole complex by dehydrating the ECM, which in turn mechanically compresses the cells (***Monnier et al., 2015***). When smaller than the exclusion size of the ECM, the osmolytes percolate through the ECM meshwork and selectively compress the cells which can then pull on the ECM (see schema in figure 1). We validate our approach by compressing ECM, cells and multicellular spheroids (MCS) using osmolytes with gyration radii *R*_*g*_ respectively larger and smaller that the ECM pore sizes (figure 2). As osmolytes, we use dextran molecules ranging from 10 to 2000 kDa. As a proxy of ECM, we use Matrigel (MG), a commercially available matrix secreted by cancer cells (***Kleinman and Martin, 2005***). To visualize the effect of compression on the ECM, we prepared microbeads composed of matrigel, with a diameter of 100 *μ*m (fig. 2a). To determine the exclusion size of matrigel, we use fluorescent dextran molecules of different gyration radii. As shown in figure 2a (top panel), fluorescent dextran molecules with a gyration radius below 5 nm (MW *<* 70 kDa, hereafter called “Small”; (***Granath, 1958***)) equally color the MG beads and the surrounding solution (left). Conversely, dextran molecules larger than 15 nm (MW *>* 500 kDa, “Big”) do not penetrate inside the MG beads, which appear darker than the surroundings (right). By following the evolution of the bead diameter (before and after 45 minutes) under a 5 kPa compression, we confirmed that small dextran molecules do not compress significantly the matrigel beads (figure 2a, middle and bottom panels). The small amount of compression, which we neglect, can be addressed by thermodynamic theories involving an interaction between the matrix and the permeating polymer (***Brochard, 1981***; ***Bastide et al., 1981***). Conversely, big dextran molecules occasion a large compression of up to 63±5% of the initial volume (quantification in figure 2b).

**Figure 1.**
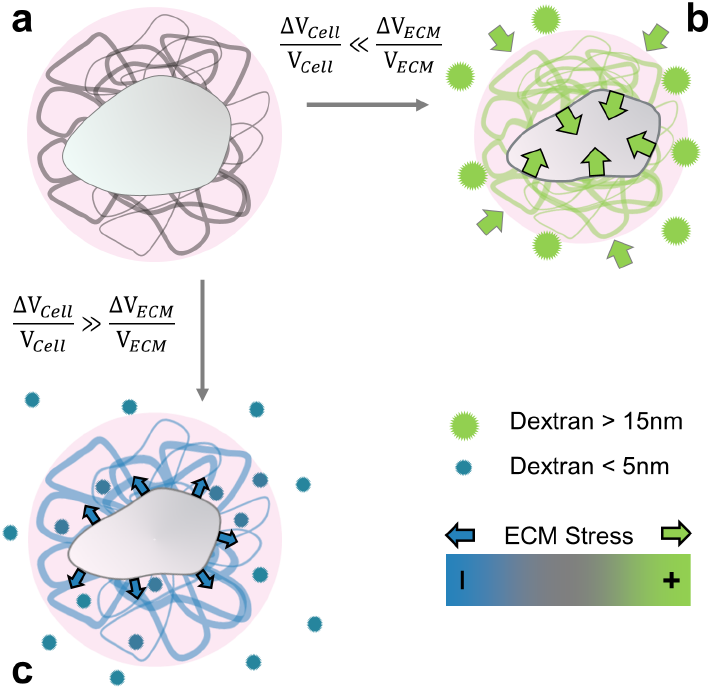
Selective compression method. **(a)** Schematic view of a cell (gray) embedded in extracellular matrix (filaments), permeated by interstitial fluid (light pink). **(b)** Big osmolytes (green) do not penetrate through the ECM and induce a global compression. Being much more compressible than the cells, the extracellular matrix absorbs most of the deformation and exert a positive stress on the cell. **(c)** Small osmolytes (blue) enter the ECM without exerting any osmotic pressure on it. Conversely, they compress the cell which, in turn, exerts a tension on the ECM.

**Figure 2.**
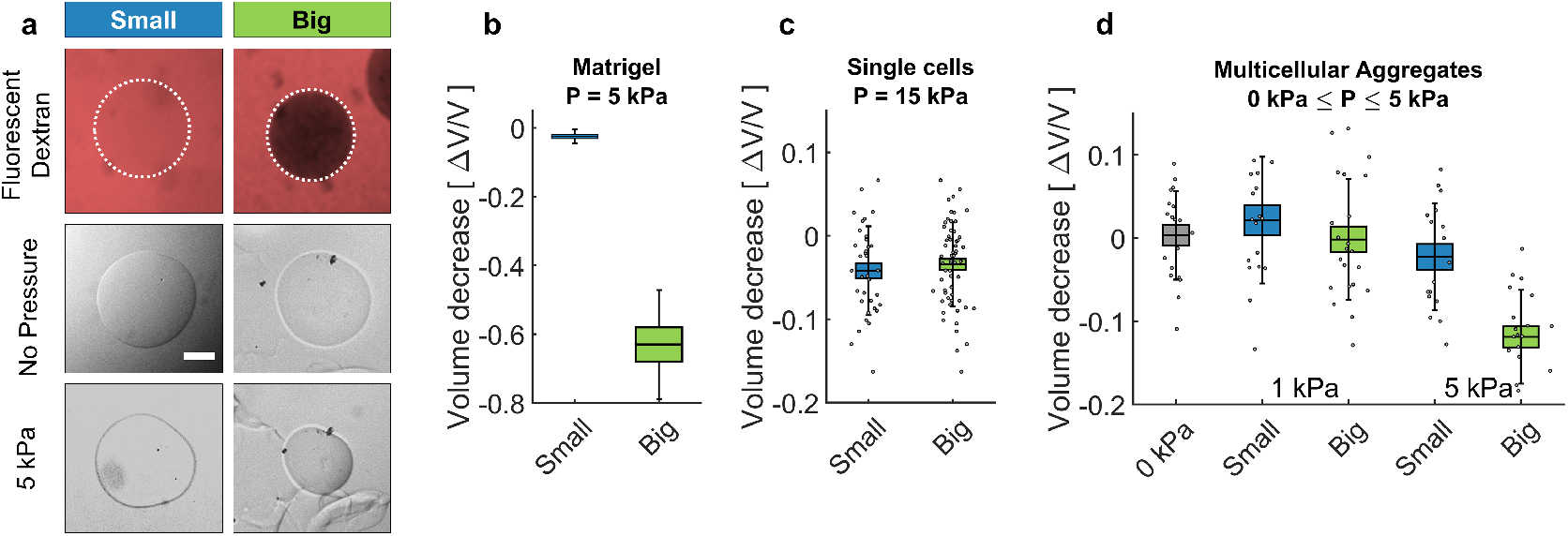
Cell and Matrigel Compression. **(a)** Fluorescently labeled dextran molecules only permeate the beads (top-left panel) only if their gyration radius is smaller than 5 nm. Otherwise (top-right panel) they are larger of the exclusion size of the matrigel network and are excluded from the bead. Compression of MG beads, occasioned by dextran molecule of two different sizes (Small: 70 kDa; Big: 500 kDa. Phase contrast images taken before and after the addition of pressure. **(b)** Beads lose 63±5% of their initial volume when compressed using big dextran, and approximately 2.5 % with small Dextran. N=10. **(c)** Compression of individual cells using dextran of different sizes, corrected for a final pressure of about 15 kPa. At 5 kPa the compressibility of individual cells is not measurable. Cell compressibility is thus negligible in comparison to that of Matrigel. **(d)** MCS relative compression under a pressure of 1 kPa and 5 kPa, exerted using small (blue) and big (green) dextran molecules. (box: ±SEOM; error bars: ±SD •: single realizations).

Analogous experiments are performed using individual cells and multicellular spheroids. Interestingly, the volume loss of individual cells is not measurable up to 10 kPa, and becomes appreciable at 15 kPa, with a relative compression Δ*V*_*c*_/*V*_*c*_ = 5 ± 5% (figure 2c). This compression indicates that CT26 cells have a bulk modulus *K*_*c*_ = 450 ± 100 kPa. In contrast to single cells, MCS are much more compressible, as they lose up to 15% of their volume under a pressure of 5 kPa (figure 2d). Furthermore, these measurements indicate that MCS have a typical bulk modulus of *K*_*s*_ ≃ 30 kPa, 15-folds smaller than that of individual cells. In contrast, small dextran molecules have no measurable effect on the volume of MCS, for moderate pressures (up to 10 kPa). However, larger pressures with these small osmolites can lead to a cell compression within the MCS associated with a swelling of the interstitial space as we show in Section 2.3.

These results confirm the ability of our method to discriminate between the effects occasioned by the compression of the whole MCS, and that due to the specific compression of the cells within the aggregate.

### 2.2 Theoretical understanding of the selective compression of composite aggregates

Our aim is to compute the displacement of the cell boundary as well as the stress applied on the cell upon the osmotic compression in both conditions. For simplicity, we consider the case of a single cell nested in a large -compared to the cell size-ball of ECM and subjected to an additional osmotic pressure Π_*d*_ (i.e. added to the existing pressure in the culture medium) obtained by supplementing the culture medium with either small or big dextran. We assume that the small dextran can freely permeate in the ECM meshwork while the big one is excluded. Our model, detailed in Section A.5, couples a classical active pump and leak model (***Hoppensteadt and Peskin, 2012***) for the cell volume regulation through ion pumping and the cytoskeleton and the ECM poro-elastic mechanics.

We show in Section A.5 that, for realistic estimates of the model parameters, both compressions with small and big dextran lead to the same cell volume loss which does not involve the mechanical properties of the ECM but only the cell volume regulation system:

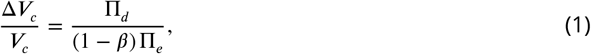

where Π_*e*_ is the osmotic pressure of ions in the culture medium and *β* ≃ 0.1 is a non-dimensional parameter representing the active pumping of ions (see Appendix A.4). For relatively low pressures (Π_*d*_ ≪ Π_*e*_ ≃ 500 kPa), the relative change of volume Δ*V*_*c*_/*V*_*c*_ is negligible. However, the mechanical stress applied by the ECM to the cell is qualitatively and quantitatively different in the two situations. For big dextran, this stress is compressive as the dominating effect of the dextran is to compress the ECM which in turn compresses the cell by the same amount. Within some realistic approximations the amount of this compressive stress (the traction force applied by the matrix on the cell) can be approximated as the applied osmotic pressure:

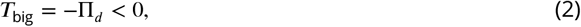

In sharp contrast with the previous situation, for small dextran, the stress applied by the ECM on the cell is tensile because the dominating effect is that small dextran does not compress the ECM but acts on the cell which compression is balanced by a tensile force in the ECM. This tension is given by

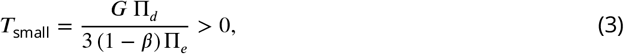

where *G* is the ECM shear modulus. Again, for a moderate osmotic shock of Π_*d*_ ≃ 5 kPa, the dextran concentration is much smaller than the characteristic ions concentration of the external medium and *T*_small_ ≤ 20Pa *≪ |T*_big_|. Therefore the presence of ECM makes the cell mechanically sensitive to a moderate osmotic compression by big dextran molecules while it is not for small dextran molecules. In both cases the cell volume is affected in the same negligible way but the mechanical stresses applied to the cell are completely different. If the osmotic compression is larger, compression with small or big dextran can induce a measurable effect on the cell volume. However, the mechanical stress applied by the ECM to the cell is different in both situations: tensile for the small dextran and compressive for the big ones.

### 2.3 Selective compression of ECM in multicellular spheroids

To test our theoretical predictions, we follow the evolution of the interstitial space inside MCS submitted to osmotic compression, occasioned either by small or big dextran. To improve the optical performance and to measure changes in the extracellular space after compression by small and big dextran, we inject the spheroids (4-5 days old) into a 2D confiner microsystem (figure 3a). MCS are confined in the microsystem for two to five hours to relax, the medium being supplemented with a fluorescent tracer to label the intercellular space. The interstitial fluorescence is measured using two-photon microscopy (figure 3b). The images of the confined multicellular aggregates are segmented with a thresholding procedure, and then the signal exceeding the threshold value is integrated over the whole aggregate to quantify the total fluorescence of the interstitial space (figure 3c). Due to optical limitations, we emphasize the effect by increasing the applied osmotic pressure to 80 kPa for small dextran and to 15 kPa for the big ones.

**Figure 3.**
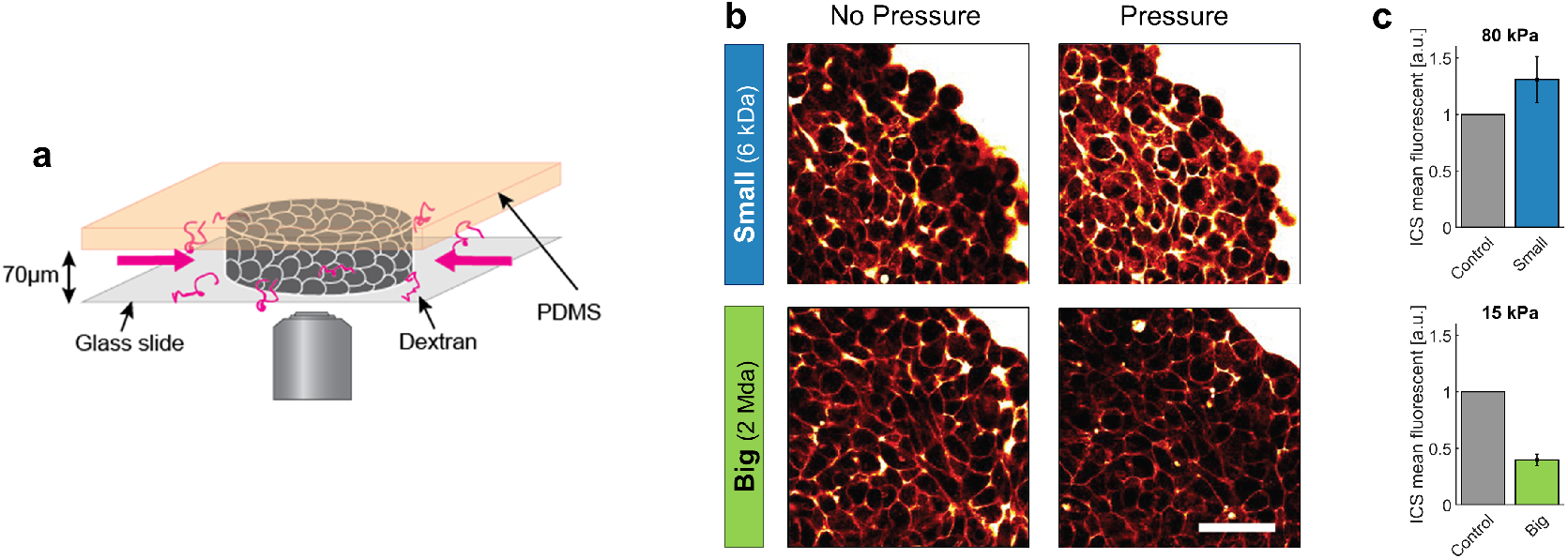
Effect of small versus big dextran on tissue intercellular space. **(a)** Scheme of the 2D confiner micro-device. The tissue is confined between the glass coverslip and the PDMS and does not move during medium exchange. **(b)**2-photon images of the tissue before and after (20min) osmotic shocks for dextran chains of 6kDa (small) and 2MDa (big), for a given mass concentration of 100g/L. Images were taken in the equatorial plane of the tissue, meaning 35*μ*m above the glass slide. **(c)** Mean fluorescence of the intercellular space averaged over the whole aggregate shown in panel B. Scale Bar: 50*μ*m

In accordance with the theoretical predictions, depending on the dextran size, we obtain two opposite behaviors. Small dextran molecules induce an increase of ~ 50% in the fluorescence intensity, measured in the interstitial space. As the total volume of the aggregate does not change significantly (figure 2d), the swelling phenomenon is related to a cell deflation compensated by the ECM swelling. In contrast, for Big dextran we measure a loss of half the fluorescence, meaning that a large amount of interstitial liquid has left the aggregate. With big osmolytes, the extracellular matrix is thus compressed as predicted by eq.(2), while the loss of cell volume is negligible compared to the latter effect. These results confirm the theoretical prediction that big and small dextran have an opposite effect on the matrix. The first puts the ECM under compression, while the latter puts the ECM under tension.

### 2.4 ECM compression controls cell proliferation and motility

To understand the role of ECM on the cell fate at longer timescale, we assess the proliferation and the motility for MCS cultured in the presence of small and big dextran. Figure 4a represents the equatorial cryosections of spheroids in the three mechanical states. Proliferating cells are labeled using antibodies against KI67. Whereas cells in control MCS (0 kPa) present a rather uniform proliferation pattern, a global compression of MCS (big Dextran) stops cell division in the core and alters the overall MCS growth, as previously reported (***Helmlinger et al., 1997***; ***Alessandri et al., 2013***; ***Montel et al., 2011***). To quantify the change of cell division rate, we monitor the volumetric growth of the spheroid for three conditions (control, small dextra, big dextran) and for several days (figure 4b). In the three cases, the spheroids initially grow exponentially (continuous lines). However, the cell division time almost doubles under compression and increases from 36±1 h (gray circles and blue squares) to 68±4 h (5 kPa, green triangles). Because experiments with MCS are typically performed in solution, where metastatic behavior is not possible, we evaluate the cell motility within the aggregate, using the Dynamic Light Scattering technique introduced by ***Brunel et al. (2020***). The mean migration velocity of cells decreases monotonously with the pressure and is reduced by 50% at 5 kPa, as compared to the unstressed case (figure 4c). Strikingly, both proliferation and motility remain almost unaltered when the MCS are exposed to an equivalent pressure (5 kPa) applied with small dextran to selectively compress the cells while leaving the native ECM unstrained (small Dextran, blue).

**Figure 4.**
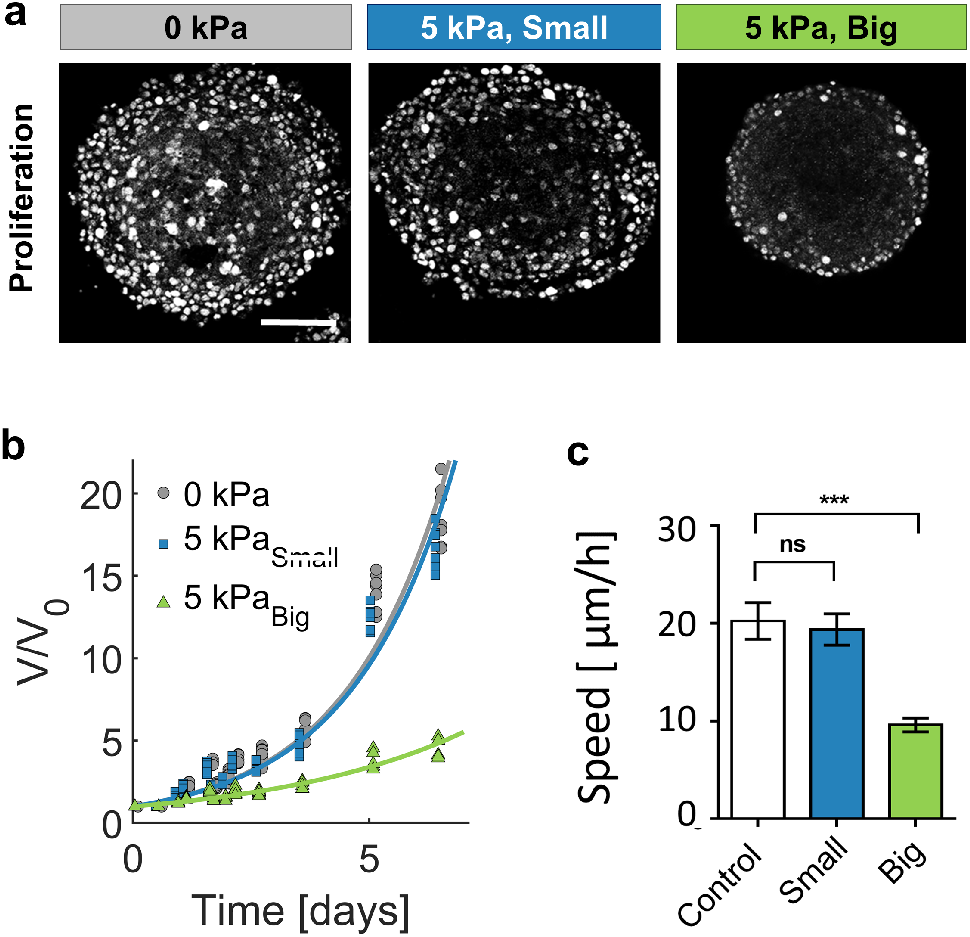
Growth of spheroids under pressure. **(a)** Proliferating cells inside MCS revealed by immunostaining of KI67 with no pressure, under overall compression of 5 kPa (big dextran) and under selective compression of the cells (small dextran). Scale bar: 100 *μm* **(b)** Volume increase of spheroids over several days, in the three reference conditions. **(d)** Cell migration speed (see SI) within MCS also significantly depends on ECM compression. N= 5 independent experiments per condition. Error bars represent ±SEM. Experiments are repeated at least on three independent samples.

Since the interstitial space is dehydrated under osmotic compression, it is possible that the cells get in contact with each other, occasioning contact inhibition of proliferation and locomotion. However, it is also possible that cells read and react to the stress accumulated in the ECM. To discriminate between these two hypothesis, we embed individual cells in a MG matrix and then compress the whole system with a 5 kPa osmotic pressure using either small or big dextran. After a few days, we observe two clearly different phenotypes. Cells that have grown without pressure are sparse in the MG, as well as cells that have proliferated in the presence of small dextran (figure 5a,left panel). Conversely, when cells are cultured in the presence of big dextran, they proliferate locally (figure 5a, right panel). Therefore, MG compression appears to inhibit cell motility and to promote the formation of mini-spheroids, which suggests that ECM compression has a direct effec on the cell-ECM biochemical signaling. The different cell morphology is particularly clear in the organization of the actin cytoskeleton. Cytoplasmic actin labeling reveals the presence of numerous protrusions, associated with high cell anisotropy in cells cultured in a relaxed MG matrix (figure 5b, left and middle panels), whereas cells appear smooth and form round structures, when the MG is compressed (figure 5b, right panel).

**Figure 5.**
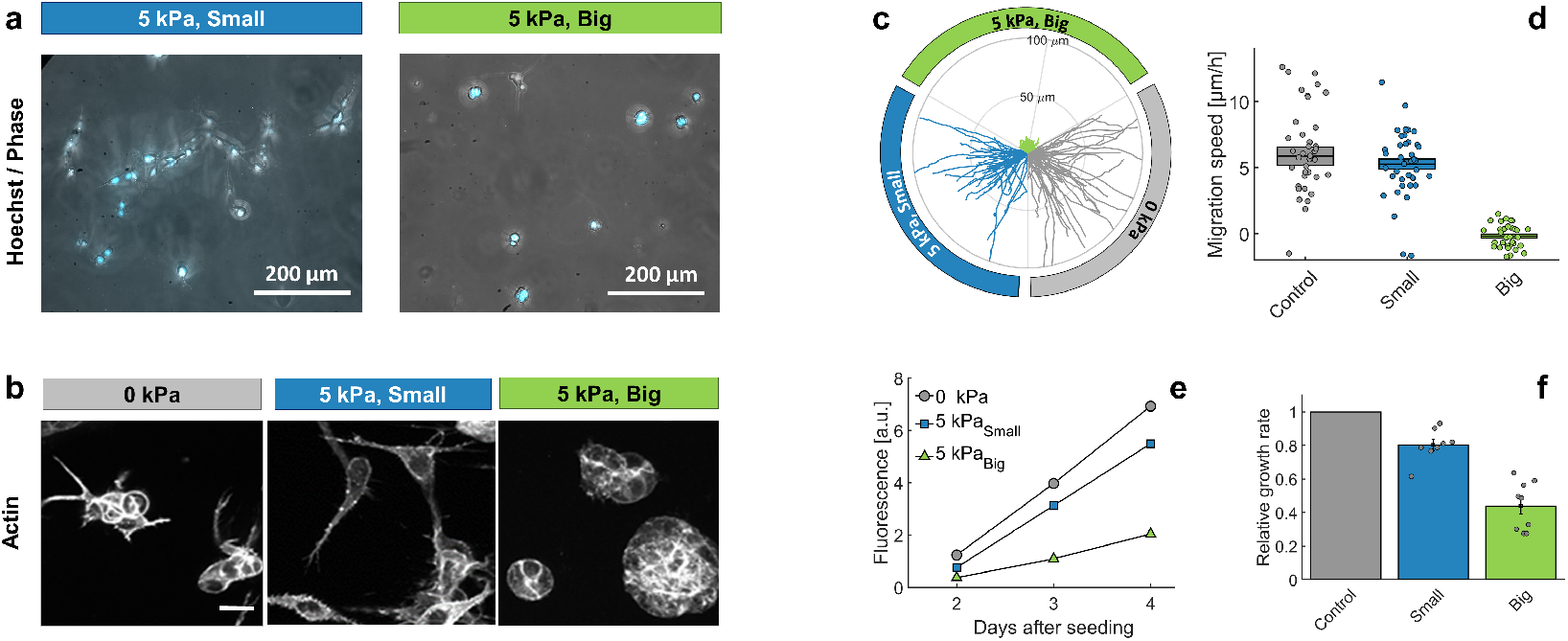
Individual cells in Matrigel. **(a)** Hoechst-labelled cell nuclei superimposed to phase image (maximum intensity z-projection). Images are taken after 2 days of proliferation in MG, either with small (left panel) or big (right panel) dextran molecules. Maximal projection from epifluoresnce stacks. **(b)** Cell morphology and anisotropy revealed by labelling of cytoplasmic actin. Maximal projection of 50 *μ*m confocal Z-stack. In relaxed MG, the cells appear more elongated and with long protrusions. **(c)** Cell motility in MG under different compression states. Starting points of trajectories are translated to the origin, to highlight the typical distance over which cells move in the three compressive states. **(d)** Quantification of in-plane velocity extracted from mean square displacements, under different compression conditions. With no pressure or with small dextran (5 kPa), the average velocities are respectively 5.8±0.8 *μm*/h and 5.2±0.5 *μm*/h. Under 5 kPa exerted by big dextran, the cells are immobile (v = 0.5±0.4 *μm*), where the error is due to tracking uncertainties. Boxes represent the average speed ±SEOM. **(e)** Temporal evolution of nuclear fluorescence intergrated over the whole sample. No pressure (o), 5 kPa with small dextran (□) and 5 kPa with big dextran (Δ). **(f)** Cell proliferation rate in the three conditions. n = 15, from 8 independent experiments.

Different morphology also correlates with different motility. Cells embedded in a compressed MG are nearly immobile, while they migrate through relaxed MG with a velocity comparable to that measured on flat surfaces. The result is summarised in figure 5c, where we report ≃40 trajectories per condition. To highlight differences and similarities between the three compression states, the starting points of all trajectories are translated to the origin and, whereas isotropic, they are compressed in three quadrants. Quantification is reported in figure 5d. From this experiment we conclude that whereas no appreciable differences are observable between the case where the MG is relaxed and the control, the cell motility dramatically drops under MG compression.

To quantify the effect of ECM compression on proliferation, we prepare several samples with the same number of cells embedded in the MG and follow the evolution of the overall fluorescence (hoechst staining) in time. Figure 5e shows the typical time evolution of the Hoechst signal in the three conditions: proliferation rate drops considerably when the MG is compressed (Big Dextran, Δ), compared to the case without pressure (o), but also compared to the case where the pressure is selectively exerted on the cells with no MG compression (Small Dextran, □). Figure 5f quantifies the mean growth rate, measured on at least 15 samples for each condition, collected on 8 independent experiments (different days and cell passages). Again, these experiments underline the impact of the ECM compressive stress on cell behavior.

## 3 Discussion and Conclusion

Large osmotic and mechanical pressures (≃100 kPa), can cause a decrease in cell volume and consequently a deformation of the cell nucleus (***Zhou et al., 2009***; ***Kim et al., 2015***) which may ultimately feedback on the cell proliferation. It has been proposed lately that the volume of cells or nucleus can be a key parameter governing crucial processes such as proliferation, invasion and differentiation ***Guo et al. (2017***); ***Han et al. (2020***). However the weak osmotic pressures (~ 1 kPa) that we apply have no measurable effect on cell volume. In addition, it is well-known from a biological standpoint that such small volume perturbations are buffered by active regulatory processes in the cell (***Hoffmann et al., 2009***). Yet, in MCS both proliferation and cell motility decrease when the aggregate undergoes a weak osmotic compression. Altogether, our results show that, for such weak compressions, the extracellular matrix (ECM) located in between the cells is directly impacted but not the cell volume. Moreover, the presence of ECM could explain the reported evidences that mechanical and osmotic pressures of same magnitudes affect the growth of MCS in the same way (***Helmlinger et al., 1997***; ***Alessandri et al., 2013***; ***Montel et al., 2011***) for osmotic pressures applied via macromolecules large enough not to penetrate the extracellular matrix. Indeed, the osmotic pressure is transduced into a mechanical one applied on the cells through the ECM drainage. As the ECM has a bulk modulus *K*_*ECM*_ ≃ 1 kPa, it can serve as sensor to measure, at the cell level, a globally applied pressure in the kPa range. Of note, stress relaxation in the ECM could occur through cleavage and remodeling of its components and such active processes should be quantified in the future.

Several mechanisms may explain how the dehydration of the extracellular matrix can result in an inhibition of proliferation and motility. First, the reduction of the interstitial space promotes interactions between neighbouring cells, which may activate contact inhibition signals of both proliferation and locomotion (***Roycroft and Mayor, 2016***). Second, the ECM porosity decreases within a compressed MCS, such that its effective permeability to oxygen, nutrients, growth factors and cytokines is reduced and might activate inhibition signals without cell-cell contact. However, both options appear incompatible with experiments of individual cells seeded in MG (Figure 5). Indeed, in these experiments, cells are isolated until the first division events. Then, when they form aggregates these have a very large surface-to-volume ratio compared to MCS, indicating that collective effects mediated by cell-cell contacts are unlikely to explain why cells decrease their proliferation. Additionally, in MCS, a key factor limiting the availability for cells of oxygen and nutrients present in the culture medium is the tortuosity of the interstitial space (***Bläßle et al., 2018***). This constraint is simply absent in experiments with single cells embedded in MG, such that the cell proliferation inhibition is most probably not related to hypoxia and starvation.

The present work therefore points at a direct mechanosensitive response of cells to the deformation accumulated in the ECM. The microscopic structure of the ECM is modified under compression (e.g. density increase, decrease of distances between cross-links and buckling of collagen fibers), with consequences on the ECM rheology. Compression is clearly accompanied by an increase in bulk modulus and, due to the fibrillar structure, to a non-trivial and non-linear evolution of the ECM stiffness (***Sopher et al., 2018***; ***Kurniawan et al., 2016***). For example, the rheological properties of synthetic ECM have been shown to affect growth of aggregates and single cells through the regulation of streched-activated channels (***Nam et al., 2019***). As integrin-dependent signals and focal adhesion assembly are regulated by the stress and strain between the cell and the ECM, the osmotic compression may steer the fate of eukaryotic cell in terms of morphology, migration and differentiation (***Pelham and Wang, 1997***; ***Choquet et al., 1997***; ***Sunyer et al., 2016***; ***Isenberg et al., 2009***; ***Butcher et al., 2009***; ***Engler et al., 2006***; ***Staunton et al., 2019***; ***Panzetta et al., 2019***). This aspect is also relevant from an oncological point of view. Indeed the ECM is strongly modified in tumour tissues and the solid stress within tumors can reach several kPa, which is in accordance with the pressure applied here (***Nia et al., 2016***). For example, there is a decrease in the ratio collagen/hyaluronan (***Voutouri et al., 2016***). The latter, more hydrophilic than the first one, tends to swell and stiffen the ECM. Whether a corrupted matrix is a contributing cause or the consequence of the neoplasia remains an open question, but the correlation between matrix mechanics and uncontrolled proliferation is more and more widely accepted (***Bissell et al., 2002***; ***Lelièvre and Bissell, 2006***; ***Broders-Bondon et al., 2018***).

In future experiments, by varying the ECM density and consequently its rheology in a controlled manner, it will be crucial to identify whether the ECM compression and the associated changes in stiffness plays a dominant role, or if - as we suggest - the mechanical stress applied on the cell through the ECM is the key ingredient directly triggering the cell biological adaptation in term of proliferation and motility.

## 4 Acknowledgements

We warmly thank J. Prost and F. Jülicher for drawing our attention to the potential impact of the poroelasticity in MCS, A. Dawid and J. Revilloud for the valuable suggestion to set the proliferation assay in MG, and C. Verdier for the valuable exchanges about the evolution of the ECM rheology under stress. This work was supported by the Agence Nationale pour la Recherche (Grant ANR-13-BSV5-0008-01), by the Institut National de la Santé et de la Recherche Médicale (Grant PC201407), by the Centre National de la Recherche Scientifique (Grant MechanoBio 2018), by the Comité de Haute-Savoie de la Ligue contre le Cancer, and by a CNRS Momentum grant (P.R.)

## 5 Methods and Materials

### 5.1 Cell culture, MCSs formation, and growth under mechanical stress

CT26 (mouse colon adenocarcinoma cells, ATCC CRL-2638; American Type Culture Collection are cultured under 37°C, 5% CO_2_ in DMEM supplemented with 10% calf serum and 1% antibiotic / antimycotic (culture medium). Spheroid are prepared on agarose cushion in 96 well plates at the concentration of 500 cell/well and centrifuged initially for 5 minutes at 800rpm to accelerate aggregation. After 2 days, Dextran (molecular mass 1, 10, 40, 70, 100, 200, 500 and 2000 kDa; Sigma-Aldrich, St. Louis, MO) is added to the culture medium to exert mechanical stress, as previously described (***Monnier et al., 2015***). To follow spheroid growth over the time, phase contrast images are taken daily. Spheroid are kept under constant pressure over observation period. Images are analysed manually using Imagej. Each experiment was repeated 3 times, with 32 individual spheroids per condition.

### 5.2 Fabrication of Matrigel beads

Matrigel beads are prepared using vortex method (***Dolega et al., 2017***). Oil phase of HFE-7500/PFPE-PEG (1.5% w/v) is cooled down to 4°C. For 400 *μ*L of oil, 100*μ*L of Matrigel is added. Solution is vortexed at full speed for 20 seconds and subsequently kept at 37°C for 20 minutes for polymerization. Beads are eventually transferred to PBS phase by washing out the surfactant phase.

### 5.3 Fluorescence eXclusion method (single cell volume measurements)

Cell volume is obtained using Fluorescence Exclusion microscopy (***Cadart et al., 2017***; ***Zlotek-Zlotkiewicz et al., 2015***). Briefly, cells are incubated in PDMS chips, with medium supplemented with a fluorescent dye that does not enter the cells. Cells thus exclude fluorescence, and one extracts cellular volume by integrating the fluorescence intensity over the whole cell. Chips for volume measurements of single cells are made by pouring a mixture (1:10) of PMDS elastomer and curing agent (Sylgard 184) onto a brass master and cured at 80oC for at least of 2 hours. Inlet and outlets were punched with a 3mm biopsy puncher. Chips are prepared few days before, bounded with oxygen plasma for 30s, warmed up at 80°C for 3 minutes then incubated with Poly-l-lysine (sigma) for 30min to 1hrs, washed with PBS, then washed with dH_2_O, dried and stored sealed with a paraffn film. The chambers are washed with PBS before cell injection. Imaging starts within 10 minutes after cell injection in order to prevent adhesion and thus cells response to the shear stress generated by the medium exchange. Acquisition is performed at 37°C in CO_2_ independent medium (Life Technologies) supplemented with 1g/L FITC dextran (10kDa, from Sigma Aldrich) on an epifluorescence microscope (Leica DMi8) with a 10x objective (NA. 0.3 from LEICA).

### 5.4 2D Confiner

The 2D confiner is microsystem conceived to image three-dimensional multicellular aggregates with a two-photon microscope (see figure 3a). Spheroids (4-5 days old) are injected in the device using a pressure controller (Fluigent, MFCS) and are partially flattened between two parallel surfaces, perpendicular to the optical axis of the microscope. Medium perfusion and exchanges is performed manually using large inlets (*>*1 mm) during two-photon acquisition. Acquisitions are performed at 37°C on a Nikon C1 two-photon microscope coupled with a femtosecond laser at 780nm with a 40x water-immersion (NA. 1.10) objective (Nikon). Chip is made by pouring PDMS elastomere and curing agent (1:10) the mold and cured for at least 2 hours. The chips are bounded to glass coverslips with 30s oxygen plasma, immediately after bounding, a solution of PLL-g-PEG at 1g/L (to check) is injected and incubated for 30 minutes in humid atmosphere to prevent cell surface adhesion during the experiment. The chips are washed with dH_2_O,dried and sealed with a paraffn film.

### 5.5 Cell culture in Matrigel

Experiments are conceived to start the culture from individual cells embedded in Matrigel. At day 1, the cells are resuspended, then dispersed in a solution containing matrigel at the final concentration of 4.5 g/l. The cells are diluted to 10,000-50,000 cells/ml, a concentration at which the average distance between neighboring cells is about 250-400 *μ*m. We therefore consider them as isolated entities. The MG/cell ensemble is gelified in 200 *μ*l wells, at 37°C, for 30 minutes. To avoid cell sedimentation, we gently flip the sample over, every two minutes. The samples are then redeposited in the incubator under three pressure conditions: no pressure and 5 kPa exerted both by small and big dextran.

### 5.6 Cells Migration in Matrigel

To quantify cell migration in Matrigel, individual cells are observed by phase contrast microscopy. Z-stacks are collected every 20 minutes and for several days, with slices spaced by 50 *μm*. Then the full stack is projected to one single layer (maximum intensity projection). Cells are tracked manually in the 2D plane, using the ImageJ MTrackJ plugin (https://imagescience.org/meijering/software/mtrackj/)

### 5.7 Cryosectioning and Immunostaining

Spheroids are fixed with 5% formalin (Sigma Aldrich, HT501128) in PBS for 30 min and washed once with PBS. For cryopreservation spheroids are exposed to sucrose at 10% (w/v) for 1 hour, 20% (w/v) for 1 hour and 30% (w/v) overnight at 4°C. Subsequently spheroids are transferred to a plastic reservoir and covered with Tisse TEK OCT (Sakura) in an isopropanol/dry ice bath. Solidified samples are brought to the cryotome (Leica CM3000) and sectioned into 15 *μ*m slices. Cut layers are deposited onto poly-L-lysine coated glass slides (Sigma) and the region of interest is delineated with DAKO pen. Samples are stored at −20°C prior immunolabelling. For fibronectin and Ki67 staining samples are permeabilized with Triton X 0.5% in TBS (Sigma T8787) for 15 minutes at RT. Nonspecific sites are blocked with 3% BSA (Bovine serum Albumin) for 1 hour. Then, samples are incubated with first antibody (Fibronectin, Sigma F7387, 1/200 and Ki67; Millipore ab9260, 1/500) overnight at 4°C. Subsequently samples are thoroughly washed with TBS three times, for 15 minutes each. A second fluorescent antibody (goat anti-mouse Cy3, Invitrogen; 1/1000) is incubated for 40 minutes along with phalloidin (1/500, Alexa Fluor 488, Thermo Fisher Scientific). After extensive washing with TBS (four washes of 15 minutes) glass cover slides are mounted on the glass slides with a Progold mounting medium overnight (Life Technologies P36965) and stored at 4°C before imaging.

### 5.8 Statistical analysis

Student’s t-test (unpaired, two tailed, equal variances) is used to calculate statistical significance as appropriate by using PRISM version 7 (graphpad Software). Statistical significance is given by *, P*<*0.05; **, P*<*0.01; ***, P*<*0.001; ****, P*<*0.0001.

## Appendix 1 A Theoretical model of the osmotic compression of a single cell nested in matrigel

Our aim is to qualitatively understand the nature of the steady state mechanical stress and displacement of a cell nested in a matrix in two paradigmatic situations:

- when some small osmolites (typically dextran) that can permeate the matrix pores are introduced in the solution,
- when some big osmolites that are excluded from the matrix are introduced in the solution.

The matrix is a meshwork of biopolymers permeated by an aquaeous solution containing ions. These ions can also permeate the cell cytoplasm via specific channels and pumps integrated in the plasmic membrane (***Hoffmann et al., 2009***; ***Lang et al., 1998***). For simplicity, we restrict our theoretical description to Na^+^, K^+^ and Cl^−^ ions which have specific channels and a well studied pump (***Therien and Blostein, 2000***) which actively pumps out three sodium ions in exchange of having two potassium ions in. Attached right under the cell membrane via some specific cross-linkers (***Diz-Muñoz et al., 2010***), the cell cortex is a thin ‘muscle-like’ actin network cross-linked by passive and contractile cross-linkers such as myosin II. The cortex has been shown to be an important regulator of the cell surface tension (***Clark and Paluch, 2011***; ***Salbreux et al., 2012***) as exemplified during motility (***Hawkins et al., 2011***) and cell morphogenesis (***Turlier et al., 2014***; ***Sedzinski et al., 2011***; ***Tinevez et al., 2009***; ***Charras et al., 2008***). The cell membrane and cortex enclose the cytoplasm a meshwork of macro-molecules permeated by water and containing the aforementioned ions. See Fig. 1 for a scheme of the model.

**Appendix 1 Figure 1.**
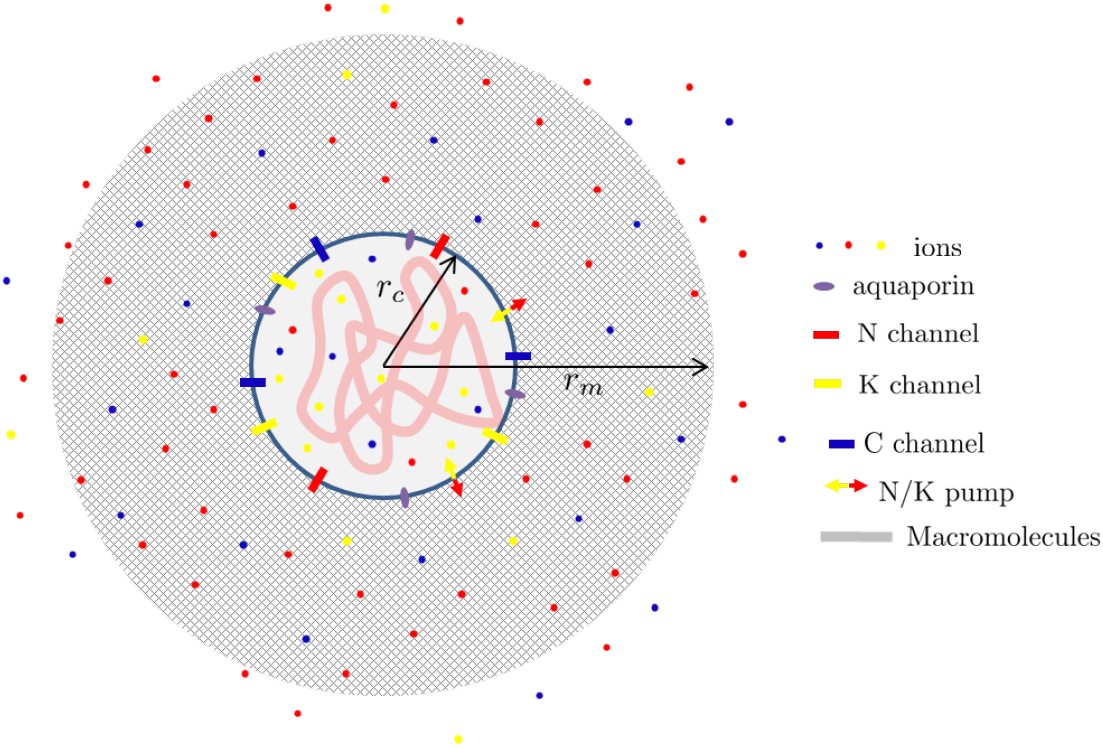
Scheme of a cell nested in a porous matrix.

For simplicity we assume a spherical geometry with a cell of radius *r*_*c*_ inside a matrix ball of radius *r*_*m*_. Each point in the space **x** can therefore be localized by its radial position **x** = *r***e**_*r*_ where **e**_*r*_ is radial unit vector. We assume a spherical symmetry of the problem such that all the introduced physical fields are independent of the angular coordinates *θ* and *φ*. Throughout this text, we restrict ourselves to a linear theory which typically holds when the deformation in the matrix is assumed to remain suffciently small.

## A.1 Conservation laws at the cell-matrix interface

## Water conservation

From Kedem-Katchalsky theory (***Staverman, 1952***; ***Kedem and Katchalsky, 1958***, ***1963***; ***Baranowski, 1991***; ***Elmoazzen et al., 2009***), assuming that the aquaeous solvent moves through specific and passive channels, the aquaporins (***Day et al., 2014***), we can express the incoming water flux **j**_*w*_ at *r* = *r*_*c*_ as (***Yi et al., 2003***; ***Hui et al., 2014***; ***Strange, 1993***; ***Hoffmann et al., 2009***; ***Mori, 2012***; ***Cadart et al., 2019***):

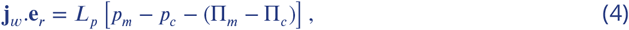

where Π_*m,c*_ denote the osmotic pressures in the matrix phase and the cell while *p*_*m,c*_ are the hydrostatic pressures defined with respect to the external (i.e. atmospheric) pressure. The so-called filtration coeffcient *L*_*p*_ is related to the permeability of aquaporins. In a dilute approximation which we again assume for simplicity, the osmotic pressure is dominated by the small molecules in solution and thus reads

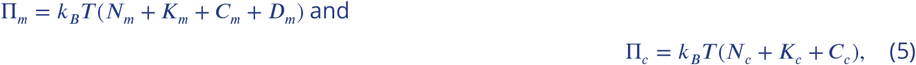

where *k*_*B*_ is the Boltzmann constant, *T* the temperature, *N*_*c,m*_, *K*_*c,m*_ and *C*_*c,m*_ are the (number) concentrations of sodium, potassium and chloride in the cytoplasm and the extra-cellular medium and *D*_*m*_ is the extra-cellular Dextran (necessary small as big are excluded) concentration in the matrix phase. We neglect in (5) the osmotic contribution associated with the large macromolecules composing the cell organelles and the cytoskeleton compared to the ionic contributions. In a similar manner, the osmotic contribution of the matrix polymer is also neglected. At steady state, the water flux vanishes (**j**_*w*_ = 0) leading to the relation at *r* = *r*_*c*_,

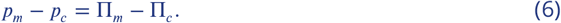

## Ions conservation

As each ion travels through the plasma membrane via specific channels and pumps, the intensities of each ionic current at *r* = *r*_*c*_ is given by Nernst-Planck laws (***Mori, 2012***),

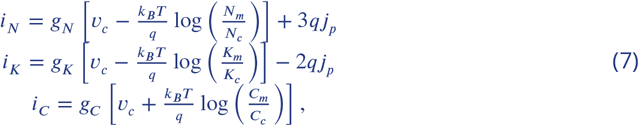

where *g*_*N,K,C*_ are the respective conductivities of ions, *V*_*c*_ is the cell membrane potential, *q* is the elementary charge and *j*_*p*_ is the pumping rate associated to the Na-K pump on the membrane which is playing a fundamental role for cellular volume control (***Hoffmann et al., 2009***). The factors 3 and 2 are related to the stochiometry of the sodium potassium pump. Again, in steady state, currents *i*_*N,K,C*_ = 0, leading to the Gibbs-Donnan equilibrium:

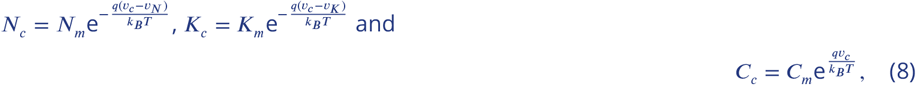

where the active potentials related to the pumping activity *v*_*N,K*_ are *v*_*N*_ = −3*qj*_*p*_/*g*_*N*_ and *v*_*K*_ = 2*qj*_*p*_/*g*_*K*_.

Supposing that the cell membrane capacitance is vanishingly small (***Mori, 2012***), we can neglect the presence of surface charges and impose an electro-neutrality constraint for the intra-cellular medium:

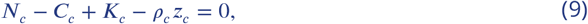

where *z*_*c*_ is the average number of (negative in the physiological pH = 7.4 conditions) electric charges carried by macromolecules inside the cell and *ρ*_*c*_ is their density. As macromolecules are trapped inside the cell membrane, we can express 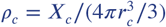 where *X*_*c*_ is the number of macro-molecules which is fixed at short timescale and only increases slowly through synthesis as the amount of dry mass doubles during the cell cycle.

## Force balance

At the interface between the cell and the matrix (*r* = *r*_*c*_), we can express the mechanical balance as

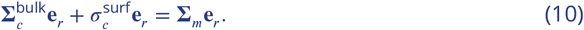

In (10), 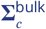 bulk is the Cauchy stress in the cytoplasm which we decompose into 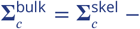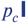, with a first contribution due to the cytoskeleton and a second contribution due to the hydrostatic pressure in the cytosol. The identity matrix is denoted **I**. The contribution due to the mechanical resistance of the cortex and membrane is denoted 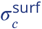. In our spherical geometry, we can express 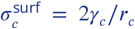 where *γ*_*c*_ is a surface tension in the cell contour. Finally **Σ**_*m*_ is the stress in the matrix phase for which we postulate a poro-elastic behavior such that, 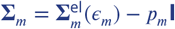 where

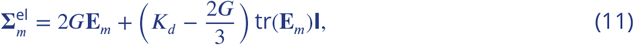

is the Hooke’s law with **E**_*m*_ the (small) elastic strain in the matrix, *G* the shear modulus and *K*_*d*_ the drained bulk modulus.

In the absence of cytoskeleton and external matrix, (10) reduces to Laplace law:

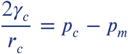

and more generally reads,

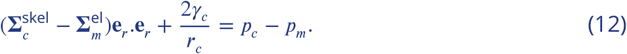

Such relation provides the hydrostatic pressure jump at the cell membrane (*r* = *r*_*c*_) entering in the osmotic balance (6) and, combining (6) and (12), we obtain

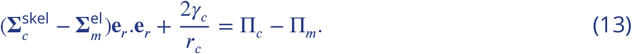

## A.2 Conservation laws in the extra-cellular matrix

## Water conservation

Assuming that the extra-cellular fluid follows a Darcy law, mass conservation of the incompressible water permeating the matrix can be expressed as

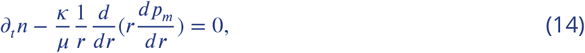

where *n* is the matrix porosity, *κ* the matrix permeability and *μ* the fluid viscosity. In steady state, *∂*_*t*_*n* = 0 and (14) is associated with no flux boundary conditions at *r*_*c*_ and *r*_*m*_ given by

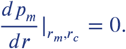

It follows that *p*_*m*_ is homogeneous in the matrix and its value is imposed by a relation similar to (6) with an infinitely permeable membrane at *r*_*m*_:

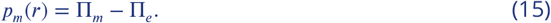

In (15), Π_*e*_ is the external osmotic pressure which reads

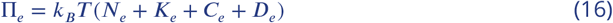

where *N*_*e*_, *K*_*e*_ and *C*_*e*_ denote the ions concentrations in the external solution and *D*_*e*_ the concentration of Dextran added to the external solution.

## Ions conservation

As we are interested about the steady-state only, the Poisson-Nernst fluxes of ions concentrations in the matrix locally vanish leading to:

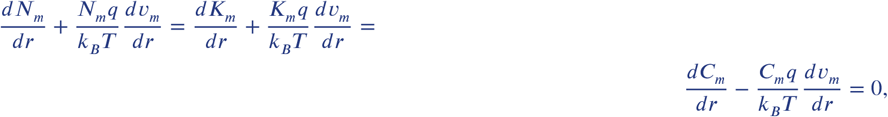

where *v*_*m*_(*r*) is the electro-static potential in the matrix.

As *v*_*m*_ is defined up to an additive constant, we chose that *v*_*m*_(*r*_*m*_) = 0 and, imposing the continuity of ions concentrations at the transition between the matrix and the external solution 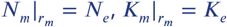 and 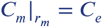, we obtain

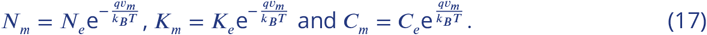

Next, we again suppose for simplicity that the capacitance of both the porous matrix and the external media are vanishingly small leading to the electro-neutrality constraints

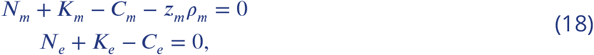

where *z*_*m*_ is the number of negative charges carried by the biopolymer chains forming the matrix and *ρ*_*m*_ is their density. As we use uncharged Dextran, its concentration does not enter in expressions (18). Using, (17) in tandem with (18), we obtain

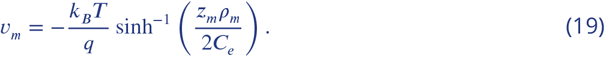

Re-injecting this expression into (17), we obtain the steady state concentrations of ions in the matrix phase:

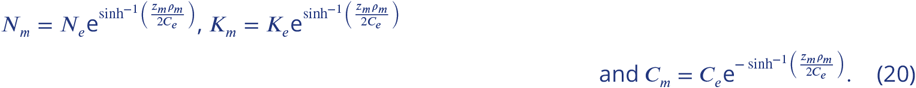

Next, we make the realistic assumption that the chloride concentration (number of ions per unit volume) is much larger than the density of fixed charges carried by the polymer chains (number charges per unit volume): *z*_*m*_*ρ*_*m*_/*C*_*e*_ ≪ 1. Indeed using the rough estimates of Section A.4, the average number of charge carried per amino-acid is 0.06 and the typical concentration of matrix is 5 g/L. As the molar mass of an amino-acid is roughly 150g/mol, we obtain that *z*_*m*_*ρ*_*m*_ ≃ 2mM while *C*_*e*_ ≃ 100mM. We can thus simplify (20) up to first order to obtain,

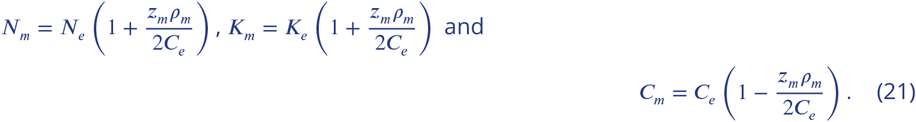

As a result, we obtain that the only steady state contribution of

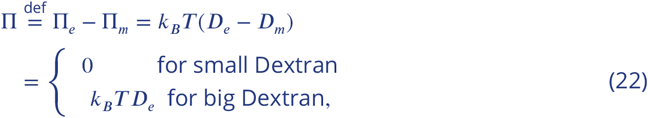

is imposed by Dextran since the ions only start to contribute to this difference at second order in the small parameter *z*_*m*_*ρ*_*m*_/*C*_*e*_. We therefore conclude that, in good approximation, Π vanishes for small Dextran molecules that can permeate the matrix and equates to the imposed and known quantity *k*_*B*_*T D*_*e*_ for big Dextran molecules that cannot enter the matrix pores.

It then follows from (15) that the hydrostatic pressure equilibrates with the imposed osmotic pressure,

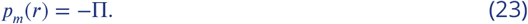

## Force balance

Using the spherical symmetry of the problem, the only non vanishing components of the stress tensor are 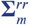 and 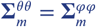. Therefore, the local stress balance reads

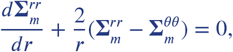

Assuming a small enough displacement, the non-vanishing components of the strain tensor are given by, 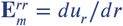 and 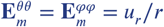 where *u*_*r*_ is the radial (and only non-vanishing) displacement component from an homogeneous reference configuration corresponding to a situation where the matrix is not subjected to any external loading and *r*_*c,m*_ = *R*_*c,m*_. Using the poro-elastic constitutive behavior (11), *u*_*r*_ satisfies

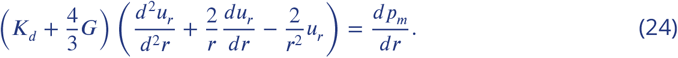

This equation is supplemented with the traction free boundary condition at *r* = *r*_*m*_

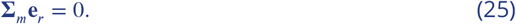

Combined with (23), the two above equations (24) and (25) lead to the solution

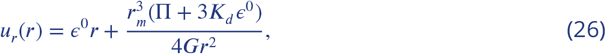

where the introduced constants *ε*^0^ is found using the displacement continuity at the cell matrix-interface:

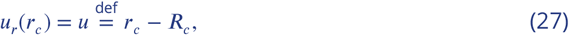

with *u* given by the change of the cell radius from a reference configuration with radius *R*_*c*_. The general expression of *u*_*r*_ therefore reads,

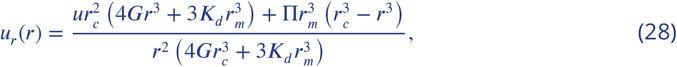

leading to the following form of the total mechanical stress in the surrounding matrix:

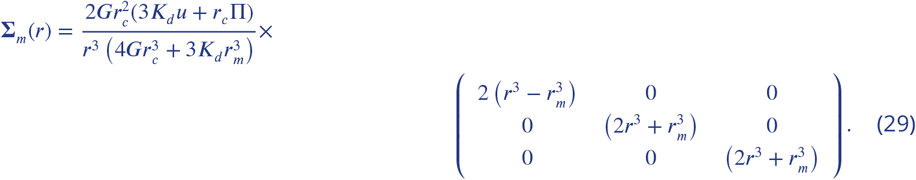

## A.3 Formulation of the model

Combining (5) with (13) and taking into account (21), we obtain the relation linking the cell mechanics and the osmotic pressures inside the cell and outside the matrix:

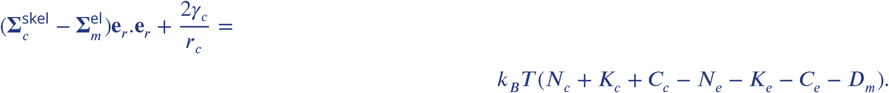

We suppose that the stress in the cytoskeleton is regulated at an homeostatic tension such that 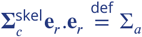 is a fixed given constant modeling the spontaneous cell contractility. We can then linearize the cell mechanical contributions close to *r*_*c*_ = *R*_*c*_ to obtain

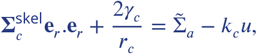

where 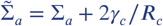 and the effective cell mechanical stiffness is 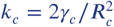.

Using and (23) and (29) close to *r*_*c,m*_ = *R*_*c,m*_ we can express,

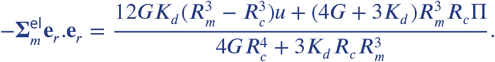

We therefore finally get the linear relation,

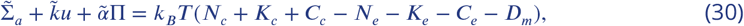

where,

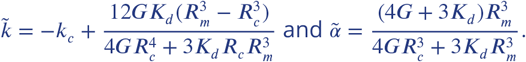

In the limit where *R*_*m*_ ≫ *R*_*c*_,

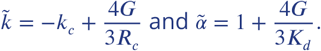

Next, using (8) and (21) and neglecting *z*_*m*_*ρ*_*m*_/*C*_*e*_ ≪ 1 we obtain the relation linking the externally controlled osmolarity with the cell and matrix mechanics:

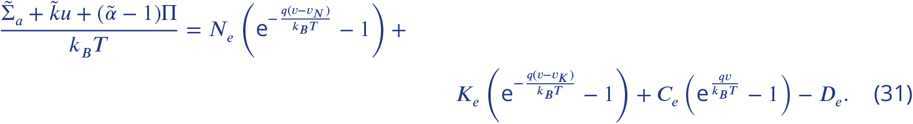

In a similar way, we combine (8) with (9) with again (21) in the limit where *z*_*m*_*ρ*_*m*_/*C*_*e*_ ≪ 1 to express the electro-neutrality condition

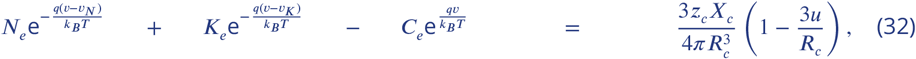

where we have additionally linearized the right handside close to *r*_*c*_ = *R*_*c*_.

The two equations (31) and (32) constitute our final model.

## A.4 Cell volume in the reference situation

We begin by computing the cell radius and the cell membrane potential in the reference configuration where by definition *u* = 0 and Π = *D*_*e*_ = 0 as no Dextran is present at all. In this case, we solve for the membrane potential 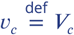 and radius *R*_*c*_ in (31) and (32) to find their reference values. This computation strictly follows (***Hoppensteadt and Peskin, 2012***).

Defining the non-dimensional parameters,

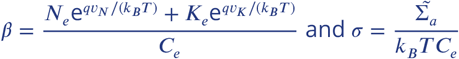

we find the reference radius and membrane potential,

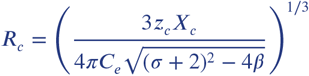

and

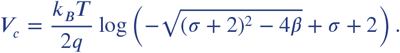

Given that the typical concentration of chloride ions outside the cell is of the order of 100 milimolar, the osmotic pressure *k*_*B*_*T C*_*e*_ is of the order 10^5^Pa (i.e. an atmosphere). In sharp contrast, the typical mechanical stresses in the cytoskeleton and the cortex are of the order of 10^2^ − 10^3^Pa (***Julicher et al., 2007***). Therefore the non-dimensional parameter *σ* is of the order of *σ* ~ 10^−3^ and will be neglected in the following. We then finally obtain the reference values,

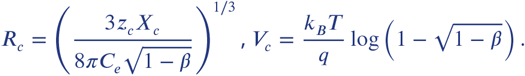

The pumping rate enables the cell to maintain a finite a volume. When *j*_*p*_ → 0, *β* → 1 and the cell swells to infinity because nothing can balance the osmotic pressure due to the macromolecules trapped inside. So it is expected that dead cells will swell and lyse. The same happens if the pumping rate is to high. Indeed as the membrane permeability of potassium is higher than the one of sodium, if the pumping rate is very high, a lot of potassium ions will be brought in (more than sodium ions will be expelled out) and to equilibrate osmolarity with the exterior, water will come in until the cell bursts because of the potassium ions pumped inside. Between these to unphysiological situations, computing the variation of volume with respect to the pumping rate, one gets that this variation vanishes when,

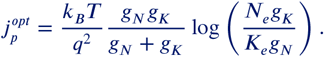

At such pumping rate the volume is less sensitive to small variations in the pumping rate that may occur.

## Rough estimates

The computation of the effective charge carried by macromolecules is complex. The folding of proteins and the electrostatic screening of charges between them (Manning effect) plays a role. See ***(Barrat and Joanny, 1997)*** for a review. We can still make a very rough estimate in the following way. We assume that macromolecules are mostly proteins. At physiological pH = 7.4, three types of amino-acids carry a positive charge, Lysine, Arginine, Histidine while two others Aspartate and Glutamate carry a negative charge. Added to this, Histidine has a pKa = 6 smaller than the pH so the ratio of [histidine neutral base]/[histidine charged acid] is 10^pH−pKa^ = 25. Hence the contribution of histidine may be neglected. The occurrence of the aforementioned amino acids in the formation of proteins is also known. The average length of proteins is roughly 400 amino acids. We subsequently obtain the average effective number of negative charges as,

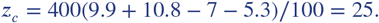

Such estimate needs to be refined and account for sugars and other macromolecules which carry more charges but a interval from *z*_*c*_ = 10 to *z*_*c*_ = 100 charges seems reasonable.

Estimate of *β* requires the knowledge of physiological external concentration of ions *C*_*e*_ = 150mM, *N*_*e*_ = 140mM and *K*_*e*_ = 10mM as well as conductances of sodium and potassium ions through the plasmic membrane. Here again the situation is complicated since the dynamical opening of channels due to some change in the membrane potential ***(Hodgkin and Huxley, 1952)*** as well as the mechanical opening mediated by membrane stretching can play a role and affect these quantities. Nevertheless a rough estimate can be given ***(Yi et al., 2003)***

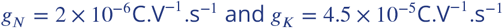

Also the pump rate is estimated in ***(Luo and Rudy, 1991***),

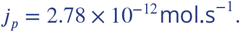

This pump rate is in good agreement with the optimal pump rate predicted by the model,

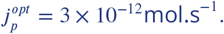

This leads to an estimate of

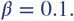

The density of macromolecules inside the cell is then found to be *ρ*_*c*_ = 3 × 10^6^ macro-molecules per *μ*m^−3^ which is a correct order of magnitude (***Milo, 2013***). To further check the soundness of the above theory we can also compute the membrane potential and obtain *V*_*c*_ = −73mV in good agreement with classical values.

## A.5 Osmotic compression of the cell

We now consider the case where, from the reference configuration presented in the previous section we impose an additional osmotic pressure in the external solution with Dextran polymers Π_*d*_ = *k*_*B*_*T D*_*e*_. We recall that according to formula (22), Π = 0 for small Dextran molecules while Π = Π_*d*_ for big Dextran molecules.

We use (31) and (32) to compute the ensuing small displacement *u*. Assuming in good approximation that the osmotic pressure imposed by chloride ions is much larger (10^5^ Pa) than the mechanical resistance of the cell cortex and the external matrix (10^3^ Pa) 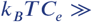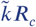 we find that,

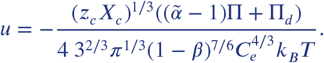

Strinkingly, making the realistic simplifying assumptions that *K*_*d*_ ≫ *G* and *R*_*m*_ ≫ *R*_*c*_, leads to the same displacement of the cell membrane in the two situations of small and big Dextran:

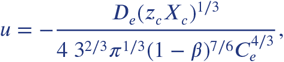

showing that the two different osmotic loading are not distinguishable at that level. The main text relation (1) is obtained by assuming that the osmotic pressure of negatively charged ions is half the osmotic pressure of all ions.

However, the mechanical stress applied of the cell is completely different in both situations. For small Dextran, the mechanical stress confining the cell reads,

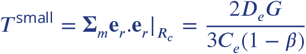

while for big Dextran it reads,

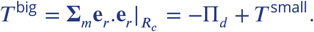

Since *T*^small^ *≪* Π_*d*_ by at least one order of magnitude, the most important feature that changes between small and big Dextran is that *T*^small^ *>* 0 while *T*^big^ *<* 0. The physical picture behind this is that small Dextran compresses the cell without draining the water out of the matrix. Therefore, the cell behaves as a small inclusion which volume is reduced by the osmotic compression. In response, the matrix is elastically pulling back to balance the stress at the interface. In contrast, for big Dextran, the water is drained out of the matrix which therefore compresses the cell.

Another important aspect is that |*T*^small^/*T*^big^| *≪* 1 so that the expansile stress applied on the cell through the matrix with small Dextran is negligible compared to the compressive stress applied on the cell by big Dextran. This is fully compatible with the idea that big Dextran induce a biological response of the cell while small Dextran is not even perceived by the cell.

Note that, like the membrane displacement, the variation of the membrane potential 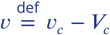 is the same in the two situations:

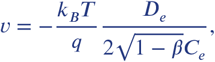

where we have made the same previous simplifying assumptions that 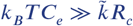, *K*_*d*_ ≫ *G* and *R*_m_ ≫ *R*_*c*_. Again such variation is negligibly small in our conditions where *D*_*e*_ ≪ *C*_*e*_ by several order of magnitudes. This further indicates that the biological response of the cell in response to a big Dextran compression has a mechanical rather than an electro-static origin.

